# DNA analysis of a large collection of shark fins from a US retail shop: species composition, global extent of trade and conservation - a Technical Report from the Monterey Bay Aquarium

**DOI:** 10.1101/433847

**Authors:** Stephen R. Palumbi, Kristin M. Robinson, Kyle S. Van Houtan, Salvador J. Jorgensen

## Abstract

We identified shark fins sampled across the entirety of a shark fin shop that had operated on the west coast of the United States until 2014. From these specimens we obtained 963 species identifications with Cytochrome oxidase (COI) sequencing and 1,720 identifications with control region (CR) sequences. We found 36-39 distinct species with COI and 38-41 with CR. Of the species identified, 16-23 are currently listed as Endangered or Vulnerable on the IUCN Red List, an additional 2 are considered data deficient, and 7 currently listed under CITES Appendix II. Of the 2.5 tonnes of fins from this collection, we estimated 56-66% (CR or COI, respectively) come from CITES-listed species or those the IUCN considers threatened or data deficient. Most of these species occur outside of the United States EEZ, comprising a global set of species that is common in most fin surveys. The principal target shark fishery in the United States (spiny dogfish; *Squalus acanthias*) has no fins in our collection. Fins seen abundantly in our collection include pelagic species such as thresher, mako, oceanic whitetip, silky, blue and hammerhead sharks, as in previous samples of the shark fin supply chain. However, in addition, we see a large flood of blacktip, dusky, sandbar, and smalltail sharks that are common in shallow coastal waters. This may indicate that the global market for shark fins takes sharks from nearshore coastal zones, all over the world. Abundant species in the fin shop included globally-distributed species such as scalloped hammerheads and shortfin mako sharks, but also regionally-restricted species such as finetooth, blacknose, and Caribbean Reef sharks found only in the western Atlantic or Caribbean. Specimens identified from rare species of particular conservation concern included the wedgefish genus *Rhyncobatus* and the white shark. Both molecular markers performed well in identifying most fins, achieving a similar degree of taxonomic certainty. The universal primers for COI regularly amplified bacteria in wet fin samples, but the CR primers were able to return shark sequences even from these degraded samples. However, the CR primers amplified a second gene, likely a pseudogene, in some important and abundant species, and seriously underestimated some species of high conversation concern such as the thresher sharks.

## INTRODUCTION

Shark fins comprise a lucrative, extensive global market that leads to the death of 70-100 million sharks a year (Worm et al., 2013), largely for the use of fin collagen (ceratotrichia) in shark fin soup. A recent study estimates that the majority of shark fin harvests are unsustainable, illegal, and that the volume traded has dramatically increased over the last decade (Sadovy de Mitcheson et al., 2018). Extensive capture of ocean-going sharks in long-line and net fisheries has led to the global collapse of many shark species (Baum et al., 2003; Nicholas K. Dulvy et al., 2008). However, the extent of the shark fin trade suggests that many other species are being similarly affected (Fields et al., 2018; Steinke et al., 2017). Recent assessments furthermore indicate some coastal shark populations have declined nearly 90% in the last 50 years (Martin et al., 2016). Efforts to monitor impacts from trade have been impeded, in part, by the extensive processing of fin products. Though some fins are traded with skin on, and with many morphological features intact, many are traded with most features removed, defying easy morphological-based cataloging.

Molecular tools have been used extensively to discover the identity of shark fin products in trade, particularly by testing fins in large wholesale markets (Clarke, Magnussen, Abercrombie, McAllister, & Shivji, 2006). Such tests have shown the widespread occurrence of ocean going sharks such as blue sharks, oceanic white tip sharks, thresher sharks, hammerhead sharks and mako sharks. But coastal species have increasingly been observed. Recently, effort has shifted to tests of retail outlets that likely integrate supply chains over months or years (Cardeñosa et al., 2018; Feitosa et al., 2018; Fields et al., 2018). For example, Fields et al. (2018) surveyed 92 retail markets in Hong Kong in 2014-2015 and processed 3,800 samples with a small piece of the COI gene, identifying them to the species level. They found 59 species and 17 other groups that could not be well identified. Cardeñosa et al. (2018) extended this survey to a total of 9,200 samples between 2014-2016 with 80% identification rate totaling 82 species or complexes. Feitosa et al. (2018) surveyed several retail fish markets in northern Brazil from 2014-2016, and from government inspections obtained in 2007, identifying 17 mostly regional and coastally-distributed species from 427 fins. In other cases, reliance on relatively small data sets [*n* =72 in (Steinke et al., 2017)] may not show the complete spectrum of commercial species. Another issue is that short DNA sequences can sometimes fail to differentiate closely related species. The balance between large sample numbers and a gene region sufficient to determine species identity has been difficult to achieve. Another opportunity is to compare retail outlets in Asia – especially at the heart of the fin trade in Hong Kong – with retail outlets in other parts of the world that have different local shark fisheries and consumer demand.

Here we test 1,720 fins collected from a retail outlet on the west coast of the US that was shuttered after shark fin sales were banned. We use sequences of the mitochondrial control region (Giles, Riginos, Naylor, & Ovenden, 2016), compared to published data sources, to estimate the identity of fins on the basis of sequence similarity and phylogenetic placement. For a subset of 961 fins, we also sequenced a region of the COI gene, giving us a comparative data set to test the ability of these two gene regions to deliver accurate identifications. Both data sets gave very similar results, showing the existence of up to 41 species in this retail collection. Detailed comparisons suggest that in some species some putative control region sequences are likely to be nuclear pseudogenes. In other samples, bacterial contamination renders COI amplification impossible but CR data appear robust.

## METHODS

### Sample collection

These data are free and open to the public, and were collected from fins provided by Jenny Giles in a collaborative research program involving Dr. Giles, Steve Palumbi, and Sal Jorgenson focused on genetic identification of shark market products at the Monterey Bay Aquarium and Stanford University (Giles et al. submitted). The samples were derived from a larger collection of fins with a range of physical conditions. Some fins occurred in a dried state, with skin attached (DR). Other dried fins had their skin removed (DRR). Some fins were more processed -wet ceratotrichia from multiple fins were pressed together in sheets (WC). Samples from disparate parts of each of these sheets were punched out and individual ceratotrichia were processed. Representative fins from each separate box or container were collected.

We used the IUCN Red List and CITES determinations to classify the conservation status of the identified species. As per previous studies, we consider species listed as data deficient to be threatened, as a central reason for being data poor is low population abundance, and most often when data deficient species are eventually classified they become threatened (N K Dulvy et al., 2014; Jenkins & Van Houtan, 2016).

### DNA sequencing and alignment

DNA extractions were performed using NucleoSpin^®^ Spin Columns (Macherey-Nagel). DNA was amplified and sequenced using standard published methods. For COI amplifications, we used modified Folmer COI primers [FishF1, FishF2, FishR1, FishR2 from (Ward, Zemlak, Innes, Last, & Hebert, 2005)], amplified with an extension temperature of 54°C, extension of 72°C, and denaturing of 94°C for 60 seconds each. For the putative control region, 374-521 bp was amplified using the primers GwF (5’-CTGCCCTTGGCTCCCAAAGC-3 ́) (Pardini et al., 2001) and 470r2 (5’- GCCATTAAAGGGAACTAGRGGA-3 ́), published in Giles et al. (2016).

Amplicons were treated with 0.75 units Exonuclease I and 1 unit Shrimp Alkaline Phosphatase (New England Biolabs) per 8 microlitre of template, and were sequenced in one direction using the forward PCR primer by Elim Biopharmaceuticals. Sequence data were aligned and ambiguities resolved manually using CodonCodeAligner ver. 7.1.2 (CodonCode Corporation, Dedham, MA). Data sets of cleaned sequences are in Appendix 1 (Master COI data file) and Appendix 2 (Master CR data file).

### Constructing COI data base

All data used in analyses were derived from public data bases. We used the 1035 sequences of the COI data base of Wong, Shivji, and Hanner (2009) to extract and download one or two individual COI sequences from each of 75 shark species. We were unable to find a COI sequence of the large tooth sawfish *Pristis microdon*, which appears in our Control Region data set (see below), although *Pristis zijsron* is available (accession EU398989.1). Likewise, two recently described mitochondrial lineages of the wedgefish genus *Rhyncobatus* (Giles et al., 2016) and the scoophead hammerhead *Sphyra media* are available from control region sequences but not COI. Our final reference COI collection consists of 177 sequences (sequences appended to Master COI data file, appendix 1).

### Constructing the CR database

All data used in analyses were derived from public databases. We found 2876 sequences from Genbank with mitochondrial control region sequences from Elasmobranchs (download Dec. 2017), including 363 full mitochondrial genomes. From this set we chose one or two sequences from each species. Because these were of variable length, we trimmed each sequence to the length between our amplification primers (Giles et al., 2016). In addition, we collected representative control region sequences from one representative product sample of each of the species that we found in our Cytochrome oxidase survey. In total our reference data base has 122 sequences from 79 species (sequences appended to Master CR data file, appendix 2).

### BLAST searches

We compared all of our market COI sequences and CR sequences to the full NCBI BLAST data base (in Jan 2018), and recorded the closest species match for each sample. Those species matches were added to the sequence sample name in FASTA format for ease in comparing phylogenetic and sequence distance results. (see appendix 1 and 2)

### Comparing COI sequences phylogenetically

We added 961 COI sequences from individual fin or fin products to the 177 sequences reference set and aligned them using the MAFFT server (Katoh & Standley, 2013). We aligned sequences with default conditions and created a Neighbor-joining phylogenetic tree using Jukes-Cantor sequence evolutionary parameters with bootstrap branch reliability scores. We inspected the resulting tree for clades containing conspecific sequences from Genbank and that contained market sequences determined to be closest to those same species by BLAST. We thus combine a phylogenetic clade-based criterion with a sequence-distance BLAST criterion for assignment of COI sequences to species. For crowded parts of the tree, particularly the genus *Carcharhinus*, we excluded all other sequences and re-constructed the tree.

### Comparing CR sequences phylogenetically

As a first step, we aligned all 1893 sequences plus 122 references using the MAFFT server. Because control region sequences evolve so quickly, there were no conserved bases in the alignment on which to build a well resolved tree. As a result, we constructed a UPGMA tree in MAFFT based on overall genetic distances. This analysis identified four major clades, allowing us to split the data set. The first clade included the lamnid sharks, and contained 423 sequences, including 346 sequences that subsequently were determined to be pseudogenes. The *Carcharhinus* sharks consisted of 1044 sequences. The genus *Spyrna* contained 234 sequences, and finally, other Carchariniform sharks consisted of 187 sequences. We re-aligned the sequences within these groups and rebuilt neighbor-joining trees with bootstrap support. As in the analysis of the COI data set, we compared the sequences within defined clades on these phylogenetic trees to their closest BLAST match in our reference database. We also cross referenced the closest BLAST match of the COI sequence from the same fin in 816 cases for which we have both data sets.

### Assigning names to sequences

We determined species names three ways. First, we determined the best match between each of our control region sequences to available databases. Second, we determined the best match of our COI sequences to these databases. Third, we examined the relationship among sequences in our collection and public reference sequences phylogenetically, looking for clusters of sequences that form species groups. Those species groups were defined by diagnostic bases, by statistical reliability of branches (bootstrap percentages), or by simple clustering. The strongest species names were defined by high confidence identification (>97% match for CR and >99% for COI) for both genes and high confidence phylogenetic clustering. For some fins we only had one DNA sequence, and so we relied on high % match for that sequence and phylogenetic clustering.

## RESULTS

Below, we detail the results of the dual analyses on COI and control region sequences. Each set of results steps through a set of phylogenetic trees that show the relationship of each sequence we collected to a reference set of sequences from public databases. The large format, full resolution trees are included as figures.

### COI sequences

In total we identified 963 fins as coming from 36-41 species. The uncertainty in species numbers is because certain pairs of species are difficult to distinguish with COI data and some species have multiple clades. Table 1 presents the summary results from COI and the control region. Also shown in this table are the sample sizes for each region, and the phylogenetic criteria used to assign species names to samples. The COI tree that we used is in a high definition pdf file in Figure 1A (all COI tree w/ bootstraps). This figure can be expanded to examine any portion of the large, complex tree.

**Table 1:** COI and CR sequences identifications for this study, compared to recently published studies.

| TAXON GROUP | FAMILY NAME | GENUS NAME | SPECIES NAME | COMMON NAME | MBA COI | MBA CR | STIENKE COI | FIELDS COI | FEITOSA COI | IUCN STATUS | CITES STATUS |
| --- | --- | --- | --- | --- | --- | --- | --- | --- | --- | --- | --- |
| thresher sharks | Alopiidae | *Alopias* | *pelagicus* | pelagic thresher shark | 38 | 8 | 12 | 19 | -- | VU | II |
| thresher sharks | Alopiidae | *Alopias* | *supercilius* | bigeye thresher shark | 170 | 95 | 8 | 37 | -- | VU | II |
| thresher sharks | Alopiidae | *Alopias* | *vulpinus* | common thresher shark | 1 | 1 | 1 | 2 | -- | VU | II |
| lamnid sharks | Lamnidae | *Isurus* | *paucus* | longfin mako shark | 12 | 11 | 3 | 4 | -- | VU | -- |
| lamnid sharks | Lamnidae | *Isurus* | *oxyrinchus* | shortfin mako shark | 168 | 306 | 8 | 133 | -- | VU | -- |
| lamnid sharks | Lamnidae | *Lamna* | *ditropis* | salmon shark | 1 | 1 | 3 | 17 | -- | LC | -- |
| lamnid sharks | Lamnidae | *Lamna* | *nasus* | porbeagle shark | -- | 0 | 5 | 6 | -- | VU | II |
| lamnid sharks | Lamnidae | *Carcharodon* | *carcharias* | white shark | 0 | 1 | -- | -- | -- | VU | II |
| ground sharks | Triakidae | *Mustelus* | *canis* | dusky smooth-hound | 1 | -- | -- | 3 | 8 | NT | -- |
| ground sharks | Triakidae | *Mustelus* | *lunulatus* | sicklefin smooth-hound | 1 | -- | 1 | 17 | -- | LC | -- |
| ground sharks | Triakidae | *Mustelus* | *mustelus* | common smooth-hound | -- | -- | -- | 10 | -- | VU | -- |
| ground sharks | Triakidae | *Mustelus* | *mosis* | Arabian smooth-hound | -- | -- | -- | 12 | -- | DD | -- |
| ground sharks | Triakidae | *Mustelus* | *higmani* | smalleye smooth-hound | 0 | 0 | 0 | 0 | 8 | LC | -- |
| requiem sharks | Carcharhinidae | *Galeorhinus* | *galeus* | school shark | 8 | -- | -- | 19 | -- | VU | -- |
| requiem sharks | Carcharhinidae | *Rhizoprionodon* | *porosus/terraenovae* | Caribbean sharpnose shark | 27 | 27 | 1 | 17 | 142 | LC | -- |
| requiem sharks | Carcharhinidae | *Rhizoprionodon* | *acutus* | milk shark | -- | -- | 3 | 66 | -- | LC | -- |
| requiem sharks | Carcharhinidae | *Rhizoprionodon* | *longurio* | Pacific sharpnose shark | -- | -- | -- | 7 | -- | DD | -- |
| requiem sharks | Carcharhinidae | *Rhizoprionodon* | *lalandii* | Brazilian sharpnose shark | 0 | 0 | 0 | 0 | 1 | DD | -- |
| carpet sharks | Ginglymostomatidae | *Ginglymostoma* | *cirratum* | nurse shark | 0 | 0 | 0 | 0 | 14 | DD | -- |
| hammerheads | Sphyrnidae | *Sphyrna* | *zygaena* | smooth hammerhead | 0 | 4 | -- | 165 | -- | VU | II |
| hammerheads | Sphyrnidae | *Sphyrna* | *mokarran* | great hammerhead | 10 | 38 | 2 | 41 | 40 | EN | II |
| hammerheads | Sphyrnidae | *Sphyrna* | *tiburo* | bonnethead | 11 | 11 | -- | 3 | 12 | LC | -- |
| hammerheads | Sphyrnidae | *Sphyrna* | *corona* | scalloped bonnethead | 11 | 6 | -- | 0.01 | -- | NT | -- |
| hammerheads | Sphyrnidae | *Sphyrna* | *lewini clade 1* | scalloped hammerhead | 10 | 24 | 7 | 0 | 18 | EN | II |
| hammerheads | Sphyrnidae | *Sphyrna* | *lewini clade 2* | scalloped hammerhead | 56 | 92 | -- | 196 | -- | EN | II |
| hammerheads | Sphyrnidae | *Sphyrna* | *media* | scoophead | 4 | 5 | -- | 0.01 | -- | DD | -- |
| hammerheads | Sphyrnidae | *Sphyrna* | *tudes* | smalleye hammerhead | 20 | 38 | -- | 0.01 | 10 | VU | -- |
| requiem sharks | Carcharhinidae | *Triaenodon* | *obesus* | whitetail shark | 4 | 2 | -- | 1 | -- | NT | -- |
| requiem sharks | Carcharhinidae | *Prionace* | *glauca* | blue shark | 42 | 72 | 7 | 1632 | -- | NT | -- |
| requiem sharks | Carcharhinidae | *Galeocerdo* | *cuvier* | tiger shark | 6 | 15 | -- | 16 | 12 | NT | -- |
| requiem sharks | Carcharhinidae | *Negaprion* | *acutidens* | sicklefin lemon shark | 3 | 5 | 1 | 29 | -- | VU | -- |
| requiem sharks | Carcharhinidae | *Negaprion* | *brevirostris* | lemon shark | 3 | 54 | -- | 5 | -- | NT | -- |
| ground sharks | Hemigaleidae | *Hemipristis* | *elongata* | snaggletooth shark | -- | 0 | -- | 5 | -- | VU | -- |
| requiem sharks | Carcharhinidae | *Lamiopsis* | *temminckii* | broadfin shark | -- | 0 | -- | 4 | -- | EN | -- |
| requiem sharks | Carcharhinidae | *Carcharhinus* | *amblyrhynchus* | grey reef shark | 8 | 34 | 1 | 15 | -- | NT | -- |
| requiem sharks | Carcharhinidae | *Carcharhinus* | *signatus* | night shark | 1 | 1 | -- | 0 | -- | VU | -- |
| requiem sharks | Carcharhinidae | *Carcharhinus* | *brevipinna* | spinner shark | 12 | 22 | 1 | 55 | -- | NT | -- |
| requiem sharks | Carcharhinidae | *Carcharhinus* | *brachyurus* | copper shark | 1 | 1 | -- | 13 | -- | NT | -- |
| requiem sharks | Carcharhinidae | *Carcharhinus* | *melanopterus I* | blacktip reef shark | 4 | 8 | -- | 2 | -- | NT | -- |
| requiem sharks | Carcharhinidae | *Carcharhinus* | *longimanus* | oceanic whitetip shark | 26 | 64 | -- | 48 | -- | VU | II |
| requiem sharks | Carcharhinidae | *Carcharhinus* | *melanopterus II* | blacktip reef shark | 3 | 0 | -- | 0 | -- | NT | -- |
| requiem sharks | Carcharhinidae | *Carcharhinus* | *leucas* | bull shark | 6 | 18 | -- | 87 | 17 | NT | -- |
| requiem sharks | Carcharhinidae | *Carcharhinus* | *perezii* | Caribbean reef shark | 1 | 20 | -- | 0.01 | -- | NT | -- |
| requiem sharks | Carcharhinidae | *Carcharhinus* | *falciformis* | silky shark | 59 | 127 | -- | 483 | 11 | NT | II |
| requiem sharks | Carcharhinidae | *Carcharhinus* | *isodon* | finetooth shark | 15 | 18 | -- | 6 | -- | LC | -- |
| requiem sharks | Carcharhinidae | *Carcharhinus* | *galapagensis=obscurus* | Galapagos/Dusky | 59 | 87 | 1 | 42 | -- | VU | -- |
| requiem sharks | Carcharhinidae | *Carcharhinus* | *acronotus* | blacknose shark | 12 | 61 | -- | 9 | 68 | NT | -- |
| requiem sharks | Carcharhinidae | *Carcharhinus* | *plumbeus=altimus* | Sandbar/Bignose | 19 | 72 | -- | 11 | -- | VU | -- |
| requiem sharks | Carcharhinidae | *Carcharhinus* | *limbatus* | blacktip shark | 97 | 254 | 1 | 219 | 9 | NT | -- |
| requiem sharks | Carcharhinidae | *Carcharhinus* | *porosus* | smalltail shark | 33 | 105 | -- | 2 | 42 | DD | -- |
| requiem sharks | Carcharhinidae | *Carcharhinus* | *amboinensis* | pigeys shark | -- | -- | -- | 54 | -- | DD | -- |
| requiem sharks | Carcharhinidae | *Carcharhinus* | *leiodon* | smoothtooth shark | -- | -- | 2 | -- | -- | VU | -- |
| requiem sharks | Carcharhinidae | *Carcharhinus* | *dussumieri* | whitecheek shark | -- | -- | -- | 14 | -- | NT | -- |
| requiem sharks | Carcharhinidae | *Carcharhinus* | *sorrah* | spot-tail shark | -- | -- | -- | 50 | -- | NT | -- |
| requiem sharks | Carcharhinidae | *Carcharhinus* | *albimarginatus* | silvertip shark | -- | -- | -- | 5 | -- | VU | -- |
| requiem sharks | Carcharhinidae | *Carcharhinus* | *macroti* | hardnose shark | -- | -- | -- | 6 | -- | NT | -- |
| requiem sharks | Carcharhinidae | *Isogomphodon* | *oxyrinchus* | daggernose shark | 0 | 0 | 0 | 0 | 14 | CR | -- |
| requiem sharks | Carcharhinidae | -- | -- | -- | 0 | 0 | -- | 114 | -- | NT | -- |
| dogfishes | Squalidae | *Squalus* | -- | dogfishes | -- | -- | -- | 19 | -- | DD | -- |
| dogfishes | Squalidae | *Squalus* | *brevirostris/megalops* | shortnose spurdog | 0 | 0 | 0 | 0 | 1 | DD | -- |
| squaliform sharks | Centrophoridae | *Deania* | *profundorum* | arrowhead dogfish | -- | -- | -- | 4 | -- | LC | -- |
| squaliform sharks | Dalatidae | *Dalatias* | *licha* | kitefin shark | -- | -- | -- | 53 | -- | NT | -- |
| chimaeras | Callorhynchidae | *Callorhynchus* | -- | elephantfish | -- | -- | -- | 27 | -- | LC | -- |
| carpet sharks | Hemiscylliidae | *Chiloscyllium* | -- | bamboo sharks | -- | -- | -- | 21 | -- | NT | -- |
| chimaeras | Chimaeridae | *Hydrolagus* | -- | chimaeras | -- | -- | -- | 14 | -- | LC | -- |
| whale shark | Rhincodontidae | *Rhincodon* | *typus* | whale shark | -- | -- | 4 | -- | -- | EN | II |
| wedgefishes | Rhinidae | *Rhynchobatus* | [ Gulf species ] | unk wedgefish | -- | 1 | -- | -- | -- | VU | -- |
| wedgefishes | Rhinidae | *Rhynchobatus* | *laevis* | smoothnose wedgefish | -- | 4 | -- | -- | -- | VU | -- |
| wedgefishes | Rhinidae | *Rhynchobatus* | *australiae* | whitespotted wedgefish | -- | 6 | -- | 26 | -- | VU | -- |
| wedgefishes | Rhinidae | *Rhynchobatus* | *djiddensis* | giant guitarfish | -- | 1 | -- | -- | -- | VU | -- |
| -- | Mixed rare | 17 rare species | -- | -- | -- | -- | -- | 26 | -- | -- | -- |

**Figure 1A:**
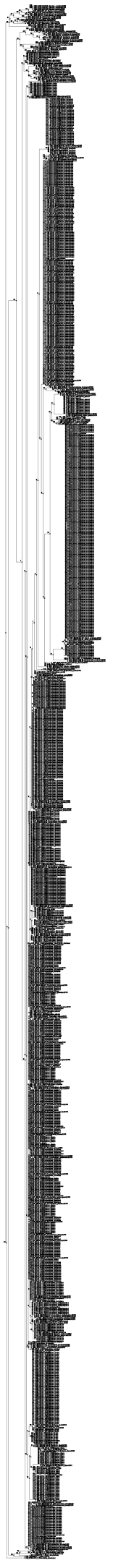
All COI tree with bootstraps. All phylogenetic trees are expandable jpg files. Each shows our market samples labeled as ‘DCA’ plus a sample number, followed by the best BLAST search of the COI sequence from that sample, where available, then the best BLAST search of the CR sequence from that sample.

**Figure 1B:**
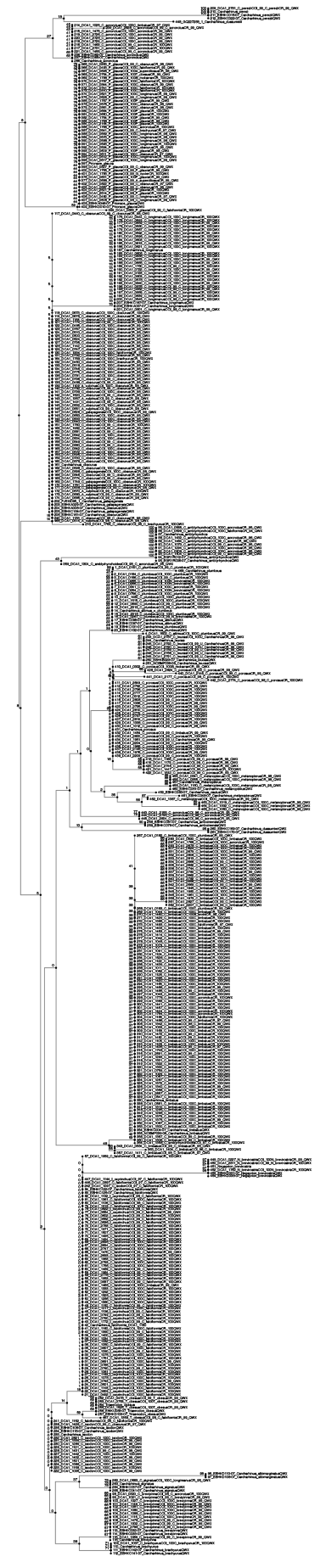
Carcharhinus COI tree with bootstraps.

A taxonomic breakdown of the COI results shows that no market fins were derived from dogfish (*Squalus*), catfish (*Apristurus*), carpet sharks (*Nebrius*), or Angel sharks (*Squatina*). Marketing of dogfish fins is less likely because of their small size. However, dogfish make up the biggest shark fishery in the U.S. and there have been calls for their fins to be exempted from state and national shark fin bans.

#### Lamniform sharks

The lamnid market samples contain all three species of thresher sharks (*Alopias*), two species of mako (*Isurus*), and the salmon shark (*Lamna ditrops*). COI clades have high bootstrap support (97%-99%) across these species, except for *A. superciliosus*, the bigeye thresher, with bootstrap support of 55% (Suppl. Fig 1A “all COI tree w bootstraps”). This species has the highest number of samples in our COI data base (n=170), closely followed by *Isurus onchyrhyncus*, the shortfin mako shark (n=168).

DCA1-0004, which has a White Shark control region sequence is not in the COI data set.

#### Carcharhinoform sharks

Genus *Carcharhinus*. The largest number of congeneric species is in the genus *Carcharhinus* and is reflected in the large number of market samples that cluster within this genus phylogenetically. Close genetic ties among these species makes some identifications with COI difficult. However, there were a number of cases where species formed well-supported clades in the bootstrap tree (Fig. 1B: *Carcharhinus* COI tree w bootstraps). The Caribbean reef shark (*C. perezii*) is a common shark in the Caribbean, and is represented here by a single COI sample in a clade defined by an 85% bootstrap. Likewise, six sequences of *C. leucas* (bull shark) are defined by a 61% bootstrap value. This *Carcharhinus* COI tree includes 14 other species clades with 41-93% bootstrap values and a total of 124 sequences (Table 1).

However, many other species of *Carcharhinus* are not well distinguishable by bootstrap phylogenetic trees because they differ by few substitutions (Wong et al., 2009). In some cases, there are diagnostic nucleotide substitutions that link sequences to certain species [(Wong et al., 2009), Table 3]. For example, our bootstrap tree shows many market samples in a large undifferentiated set representing reference sequences for *C. falciformis, isodon, galapagensis and obscurus*. Yet, *C. falciformis* and *C. isodon* each have two diagnostic bases in the COI region, and *C. galapagensis* can be distinguished from *C. obscurus* at two others. We used these diagnostic SNPs (listed in our Table 1) to provide species names and could then compare these names to the clades in the phylogenetic analysis. As a result, a UPGMA tree shows a *C. falciformis* clade that corresponds largely to the 59 sequences that contain the C*. falciformis* diagnostic SNPs. Likewise, a *C. isodon* clade in the UPGMA tree corresponds to the 15 sequences that show the diagnostic *C. isodon* SNPs. Diagnostic SNPs in COI do not appear to exist to distinguish *C. galapagensis* and *C. obscurus* (Wong et al., 2009). However, two SNPs define the COI clade that contains these two species and in our COI data set, we observe 59 sequences. Another problematic species mixture is the sandbar and bignose sharks, *C. plumbeus* and *C. altimus*. These cannot be distinguished by COI sequence alone and make up 17 sequences in our COI data base. For *C. acronotus*, we find two distinct sequence clades that BLAST to this species at 96% or 100% identity (n=3, n=9, respectively). Whether the three sequences with lower identity in a separate clade represent a different species will require further research. The widespread blacktip reef shark (*Carcharhinus limbatus*) forms a UPGMA clade with 97 sequences. Less widely distributed is the smalltail shark (*C. porosus*), found from the Gulf of Mexico to northern Brazil. Despite its small size and restricted range, we found 33 sequences.

#### Hammerhead sharks

The hammerhead clade within the Carcharhinoformes is represented in our data set by six of the eight named species. The local endemic *Sphyrna gilberti* (Quattro, Driggers III, Grady, Ulrich, & MA, 2013) is missing, possibly because there is no recorded COI sequence. *S. zygaena* is also missing from our COI data, although this species occurs in the CR data set. *S. lewini* occurs in three separate clades. Clade I, with ten sequences of *Sphyrna lewini* appears outside the main *Sphyrna* clade in the bootstrap phylogeny. This group contains sequences only 96% identical to a second clade of *S. lewini* that occurs more deeply imbedded in the *Sphyrna* clade. These two intraspecific clades have the same COI amino acid sequence and differ at only silent sites. This group is also 100% identical to a number of *S. lewini* sequences on Genbank. The second *lewini* clade has the most hammerhead sequences in our sample (n=56) and also has several voucher reference sequences on Genbank with 100% identity. These two clades may represent different species, but we will count them together in our data summary.

The third *S. lewini* clade has no Genbank or BOLD reference sequences. Our sequences that fall into this clade have 11 CR sequences that BLAST to *S. corona* rather than to *S. lewini*. As a result, this third clade, is probably *S. corona*.

#### Other Carcharhiniform sharks

Within the family Carcharhinidae, but outside the genus *Carcharhinus*, several genera have defined clades and are found in our collection. These include the tiger shark (*Galeocerdo cuvier*, n=6), and two species of lemon sharks (*Negaprion*, n=6). *Rhizopriodon* species (sharpnose sharks) are extremely similar in their COI sequences and our phylogenetic approach cannot distinguish *R. porosus* from *terraenovae* for these 27 sequences. By contrast, *R. lalandii* and *R. acutus* appear to have slightly different COI sequences but do not appear in the COI data set.

Outside the family Carcharhinidae, there are market samples from a number of Carchariniform sharks in different families. The school shark, *Galeorhinus galeus* occurs in a small clade of eight samples. Batch BLAST searches originally identified our *G. galeus* sequences as coming from *C. dussumieri* or *C. obscurus*. However, a manual recheck shows 99-100% identity with the Wong et al. (2009) sequences of *G. galeus*. Two market sequences were included in the clade that included the genus *Mustelus*. A manual BLAST shows this sample to be 100% identical to *M. lunulatus*. The other sequence was named by BLAST as *M. canis*, and is 100% identical to other *M. canis* in Genbank.

#### Poor sequence reads

In addition to these 969 full length, high quality sequences, we obtained 149 sequences with poorer read quality or shorter length. Phylogenetic analysis of these sequences using the same Genbank reference libraries reveals a similar number and distribution of species as the higher quality sequences. In particular, no species were seen in this set that had strong placement in species clades not included in the previous analyses. Because these phylogenetic placements were in general more equivocal than for the high quality sequence, we do not include them in our data summary. In addition, we produced 287 COI sequence reads that were such poor quality that they had lengths under 300 bp (n=144) or between 300 and 500 bp (n=143). These sequences were not analyzed further.

#### Bacterial amplification of shark products

The final category of COI sequences is a set of 369 amplification products from fin samples that BLAST to bacteria. Of these 244 are identified as *Pseudomonas fluoresens*, *P. sp*. or *P. veronii* (n=62, 174, 8 respectively). The rest are identified as generic bacteria, or in the genera *Kangiella, Oceanimonas, Ensifer*, or *Vibrio*. Poor sequence quality reduced BLAST confidence in 97 sequences. Most of these bacterial sequences (311 out of 369) are derived from wet ceratotrichia packaged to allow rapid use in restaurants.

### CR Sequences

BLAST searches of our CR sequences on Genbank find a >95% match to publicly available data for 1,528 of our sequences. Embedded in that number are 1,496 sequences with a >97% match. Left behind are 346 without a reputable >95% match. The largest part of that group is a set of sequences that show a maximum of 77-80% match to any control region sequence in the publicly available control region data for sharks. Below we suggest that these are pseudogenes and conduct a separate analysis to ascertain their value in fin identification.

For each clade in each of the phylogenetic trees (see methods), we counted the number of fin CR sequences for which there was the same closest species match for CR and COI (CR=COI), the number for which the two sequences did not match (CR<>COI), the number of samples for which we did not have COI sequence (No COI), and the number of reference sequences from the database. In total, we counted 1888 CR sequences, of which 496 matched the COI sequence, 320 did not match, and 1072 had no COI sequence. The breakdown of these sequence matches by species is in Table 2.

**Table 2:**
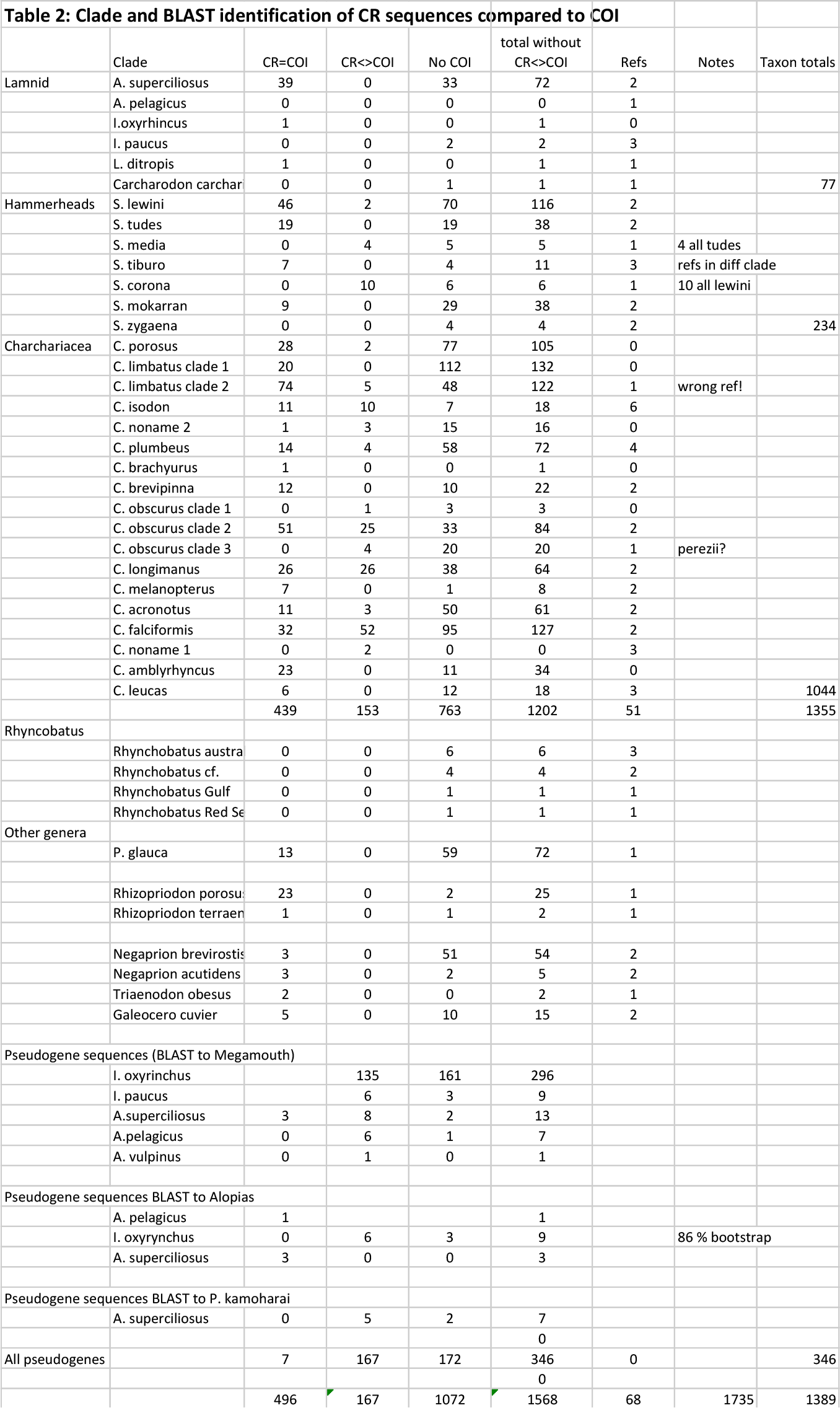
Comparison of best BLAST or phylogenetic identification for CR and COI gene sequences. CR=COI represents the number of fins that showed the same identification for both genes. CR<>COI represents fins with identifications that do not match. No COI represents fins for which there is no COI sequence. Refs is the number of reference sequences from Genbank in the phylogeneytic clades defining species.

#### Genera outside major clades

There are no sequences in our CR data set from the dogfish *Squalus*, or any sequences in this family, a conclusion that matches our COI data set. In the wedgefish genus *Rhyncobatus*, we find four species with high bootstrap support (N=6,4,1,1 respectively). This genus is undergoing taxonomic revision (e.g. Giles et al. 2016) and the names of two of these species are not yet determined. The two fins that are identified in the genus *Mustelus* in the COI data set were not sequenced as part of the CR data set, and this genus does not appear in any other sample.

#### Carcharhiniformes

The large number of insertions and deletions in the CR data set interferes with bootstrap analysis because there are few conserved sequence positions that can be included. However, a bootstrap tree shows distinct clades including sequences identified by Genbank samples as the blue shark *Prionace glauca* (bootstrap 100%, n=72), *Rhizoprionodon porosus* (bootstrap 89%, n=25), *Negaprion acutidens* and *N. brevirostris* (bootstrap 49%, n=5,54, resp.), *Galeocerdo cuvier* (bootstrap 100%, n=15), and *Triaenodon obesus* (bootstrap 100%, n=3). (See Fig 2. “Other genera CR tree”).

**Figure 2:**
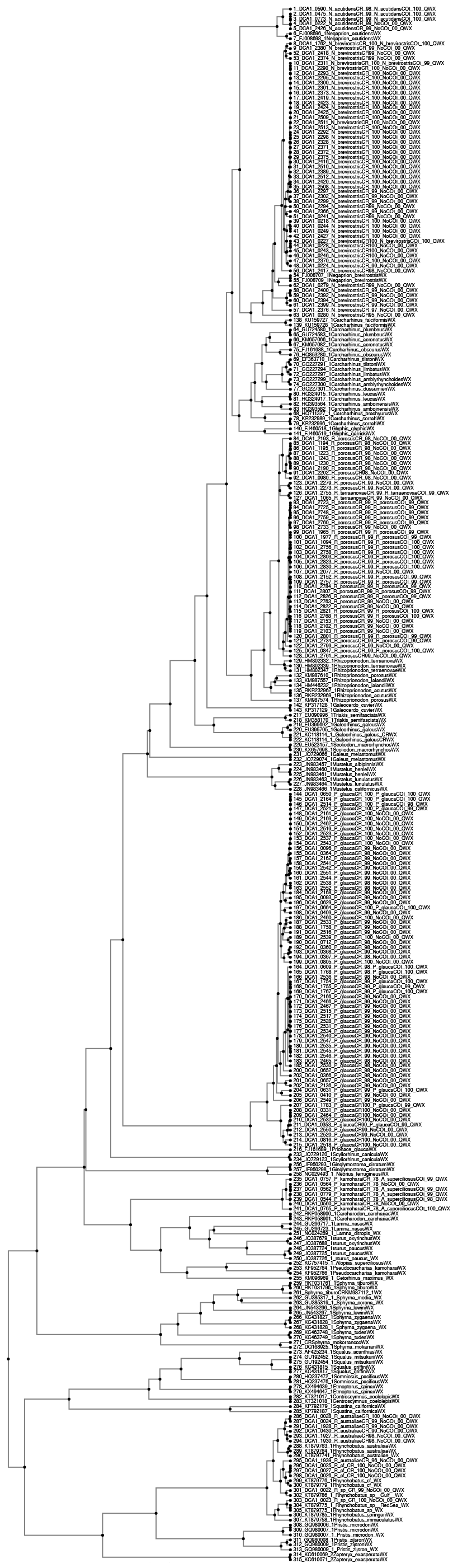
Other genera CR tree

In addition, there are a large number of sequences that cluster in the genus *Carcharhinus*. To improve tree resolution, these were grouped separately into a separate sequence file trimmed to 475 bases to match reference sequences, and run in MAFFT to produce a large UPGMA tree (Fig. 3 Carcharhinus CR tree). This improved tree still shows poor bootstrap resolution among most *Carcharhinus* CR sequences. Nevertheless, within this tree, a series of clades defines a large number of species and sequences. We assign a species name to fin sequences based on their inclusion in distinct sequence clades named by reference samples from Genbank. Further confidence in these clades and assignments comes from evaluating these assignments based on whether the CR identification matches the COI identification. In a few cases, no Genbank sequence exists, and we assign clade names based on COI assignments from the same fins. The following summarizes these results from the bottom to the top of the tree.

**Figure 3:**
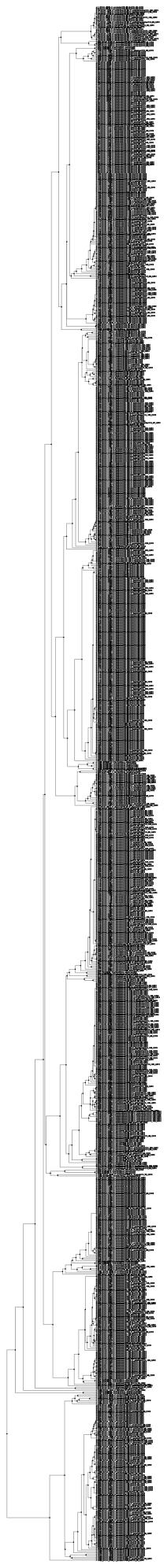
Carcharhinus CR tree

*C. porosus*, the blacktip shark, occurs in several large clades of similar sequences (ca. 99% identical, n =107 total). *C. limbatus* is the most abundant species and occurs in two major clades (n =132, 127 respectively). Two representative sequences from those two clades DCA1_0172 and DCA1_0173 are 98.6% identical, differing at 4 transitions and 2 transversions. By contrast, *C. isodon* has one small clade of sequences (n=28), highly similar to control regions from whole mitochondrial genomes for this species. A group of 19 sequences cluster together but show highest BLAST hits in the 96-97% identity range for a variety of *Carcharhinus* species. The COI sequences of five of these also show similarity to a large number of different species, never the same as the best CR BLAST hit. It is not possible to determine what species these 19 sequences derive from these data.

For COI, *C. plumbeus* and *C. altimus* form a group that cannot be easily distinguished. Although there are CR sequences for *C. plumbeus* in public data bases, there are currently no *C. altimus* CR sequences. In our CR data set, *C. plumbeus* sequences make up a large clade (n=76). Two of the sequences form a clade within this clade and have high affinity to COI *C. altimus* data (100%) and lower affinity to *C. plumbeus* CR data (98-99%). Whether these two sequences represent *C. altimus* (and the others represent *C. plumbeus)* will demand a series of positively identified individuals sequenced for the control region.

A single sequence of *C. brachyurus* occurs in our data set, clusters well with the reference sequence and has a COI sequence highly similar (100%) to *C. brachyurus*. In this clade is also a subclade of *C. brevipinna* with 22 sequences. A large clade of sequences of *C. obscurus* occurs in three subclades. Subclade 1 has four sequences with high similarity to the reference *C. obscurus* whole mt genome. A second, large clade shows 98-99%% similarity to the first clade, with 109 sequences. A third clade is a mixture of sequences with highest affinity to *C. obscurus* (n=16) or *C. perezii* (n=8). One sequence in this third clade is 99% identical to *C. perezii:* this same fin has a 99% COI identity to *C. perezii*. Whether this clade of 24 sequences actually represents *C. perezii* might require further sampling. The species *C. longimanus* shows a single clade with 90 sequences, whereas *C. melanopterus* occurs in a clade of 8. A clade of *C. acronotus* has 64 sequences, and *C. leucas* appears with 18. The second most abundant species is *C. falciformus* (n=179) which occurs in a single clade in our data set.

Last, there is a clade of 34 sequences that show no more than 94-96% identity to the closest control region sequences in Genbank. Of the 12 fins with matching COI sequence, there is 100% identity to *C. amblyrhyncos* in nine of them. (The three other fins have *Alopias* COI sequences.) There is no *C. amblyrhynchos* CR sequence on Genbank, but we provisionally ascribe these 23 sequences to *C. amblyrhynchos* based on their match to COI.

#### Hammerhead clade

The hammerhead group is represented by 234 sequences from the same species and clades as the COI data set with the addition of four sequences from *Sphyrna zygaena*. Unlike the Carcharinids, bootstrap values for hammerhead species assignments were high – ranging from 70-100% (Table 1). As in the COI data set, there were two separate clades of control region sequences in *S. lewini* and a separate clade for *S. corona*. There were 38 sequences of *S. mokarran*, two of which appeared in a related but 3-4% divergent clade. (Fig 4: Sphyrna CR tree).

**Figure 4:**
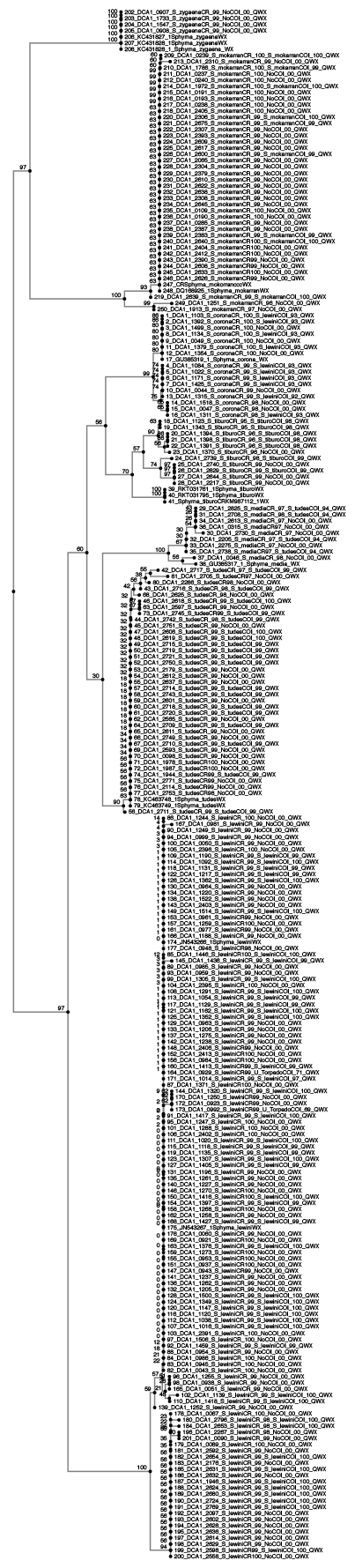
Sphyrna CR tree.

#### Pseudogene sequences in Lamnid sharks

Our phylogenetic data and BLAST searches for the CR sequences show a virtual absence of *Isurus* or *A. pelagicus*, despite the large occurrence of these taxa in the COI data set. Many CR sequences from fins identified with COI in *Isurus* and *Alopias* are a poor match to vouchered CR sequences. For example, sample DCA1-0745 has an *Alopias pelagicus* COI sequence (100% identical to JN315429.1), but its CR sequence is only 75% identical to the control region sequence from the *A. pelagicus* mtDNA genome sequence on Genbank (NC_022822.1). The closest match to the CR sequence of DCA1-0745 is to a Lamnid shark in the genus *Megachasma*, at 77% identity. Overall, we have 299 sequences that BLAST best to this genus at 77-81% identity.

To investigate this further, we generated a phylogenetic tree from control region sequences of Lamnid species from Genbank including *Isurus* and *Alopias* (Fig. 5). Genbank sequences from full mitochondrial genomes group together in this clade, along with sequences from other lamnid sharks including two *Isurus* species, *A. pelagicus* and a large number of our market sequences from *A. superciliosus*. (red labels in Figure 5). However, most of our market sample CR sequences from lamnid sharks form a clade separate from the clade defined by vouchered whole mitochondrial genomes. (upper, black labelled samples in Fig. 5). The other clade differs by about 20-24% and has no voucher sequences from full mitochondrial genomes. Instead it has exclusively sequences we generated with our CR primers from market samples. These include most of our market sequences from *I. onchyrhyncus*, *I. paucus*, and *Alopias pelagicus* samples.

**Figure 5:**
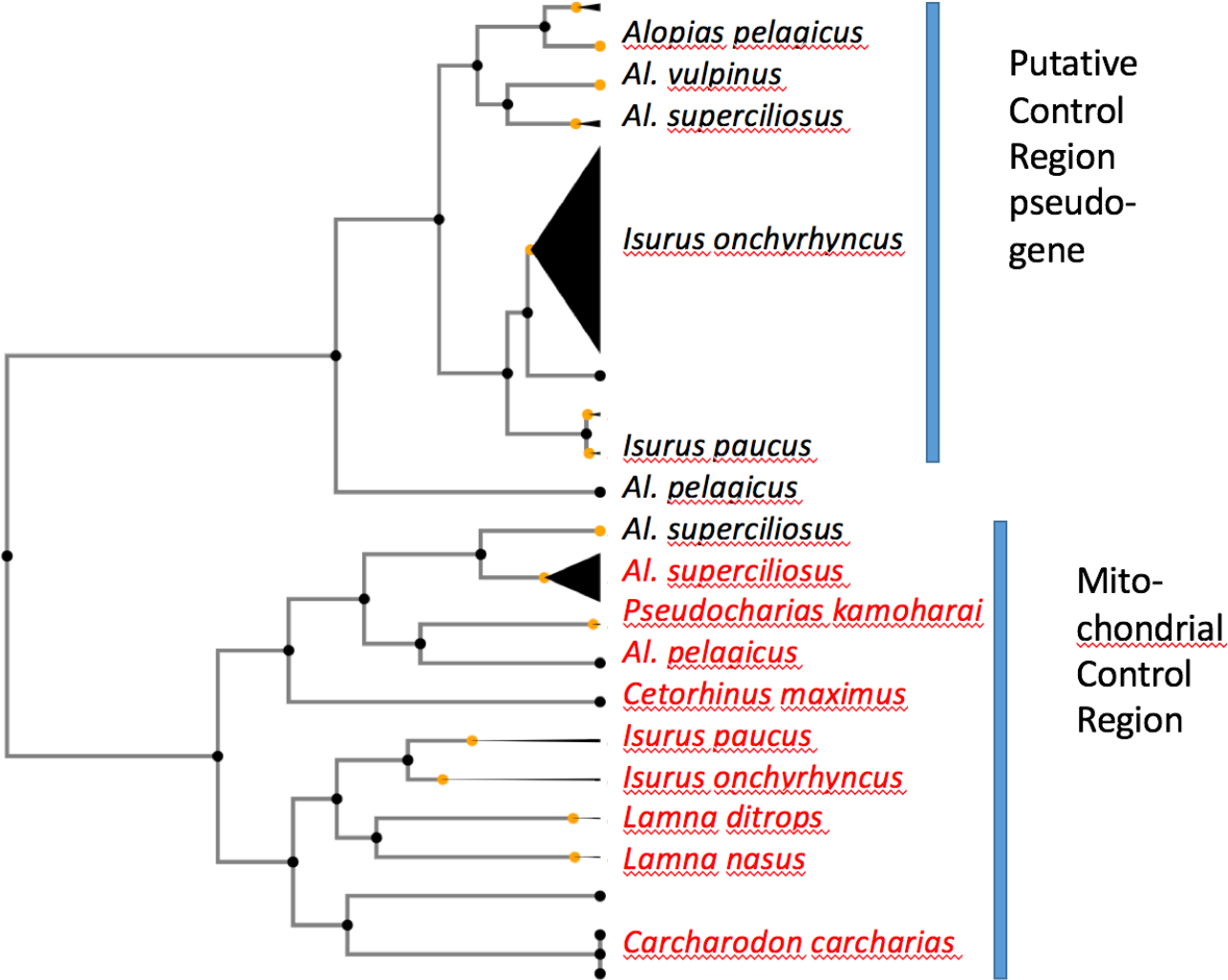
Unexpected amplification of pseudogenes. The figure shows phylogenetic relationships among mitochondrial control region sequences (red labels) from Genbank and sequences from MBAq fins derived from our control region primers (black labels). The top black-labelled cluster of sequences from *Alopias* and *Isurus* species are 20-25% divergent from known mitochondrial sequences from the same species. These black-labelled sequences are probably mitochondrial pseudogenes, sequences originally from the mitochondria that were passed into the nucleus long ago and no longer function. One exception is a set of *Alopias superciliosus* sequences from our data set that appear to be valid control region sequences.

One overall explanation for these data is that our control region primers typically amplify a different, but related, gene region in *Isurus* and most *Alopias* sharks. Because there are no duplicate control region sequences in the *Alopias* full mt genome, this duplicate, related gene region is likely to be a mitochondrial pseudogene (e.g., Bensasson, Zhang, Hartl, & Hewitt, 2001).

#### Species differences show lower divergence among pseudogenes

The clade of sequences from validated mitochondrial genomes differ between species more than do our amplified fragments from the other clade. For example, nucleotide divergence of *I. paucus* and *I. oxyrhincus* is 8% for Genbank mitochondrial CR sequences: transversions outnumber transitions 19 to 12. Between the *I. paucus* and *I. oxyrhinucs* sequences we amplified in the other clade, we find only 1.5% divergence (4 transitions and 3 transversions) along with 18 insertion/deletions. High divergence in the validated mitochondrial clade is expected given the generally higher rate of molecular evolution in mtDNA (Avise, 2012). By comparison, COI sequences from these same two *I. paucus* and *I. oxyrhinucs* individuals differed at 12% of 631 sites, including 58 transitions and 17 transversions, suggesting that the lower sequence clade in Figure 6 is evolving much like other mtDNA genes.

**Figure 6:**
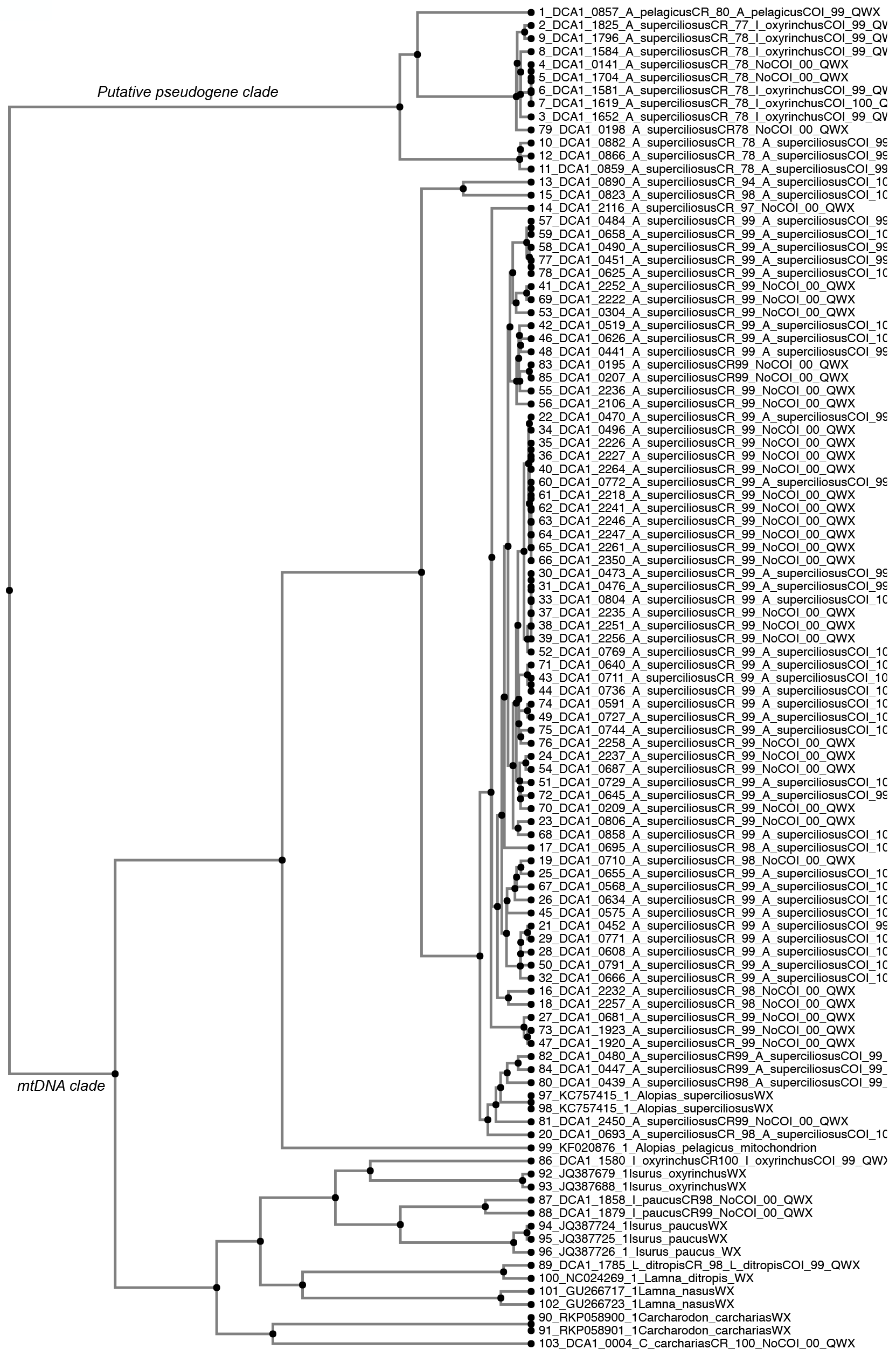
Lamnid CR sequence tree. Sequences that amplify with CR primer and that cluster with CR sequences from whole mitochondrial genomes are shown in the lower clade, suggesting that these sequences are from true Control Regions section of mtDNA. The upper clade, starting with DCA_0857 are CR amplified sequences that are highly divergent, suggesting they are pseudogene sequences.

#### True Lamnid mtDNA CR sequences in some species

The genus *Isurus* comprises a sizeable fraction of our COI data set, but there are only three sequences from our CR data that BLAST highly and cluster phylogenetically to the available *Isurus* CR sequences. Likewise, *Alopias pelagicus* is common in our COI data set but absent in the CR data set despite a reference control region sequence from the complete mitochondrial genome of this species. By contrast, we have 72 sequences of *Alopias superciliosus* that form a good clade with the control region sequence from the mitochondrial genome of this species (lower clade in Figure 6). In addition there are sequences from *I. oxyrhincus* (n =1), and *I. paucus* (n=2) in the clade of true CR sequences.

#### Two genes amplify from CR primers in some species

Multiple sequence products from individual fins can complicate identification. In our case, we sometimes find sequences from different individuals of the same species in different clades: for example, sequences from two *I. oxyrhincus* individuals identified as conspecifics by COI sequence appear in different CR clades and differ by 21%. Likewise, phylogenetic trees of *Alopias superciliosus* CR sequences from our fin sample suggest a mixture of valid control region sequences and pseudo-CR data (Fig. 6, versus Fig. 7). Altogether, the phylogenetic tree shows 72 sequences of *A. superciliosus* CR and 11 pseudoCR. The other two species, *A. vulpinus* and *A. pelagicus* have 1 and 7 pseudoCR sequences, respectively. By contrast, *Isurus oxyrhincus* shows a large number of pseudoCR sequences (296) and perhaps one true CR. The species *I. paucus* also is dominated by pseudoCR sequences (n=9), with two valid CR.

**Figure 7:**
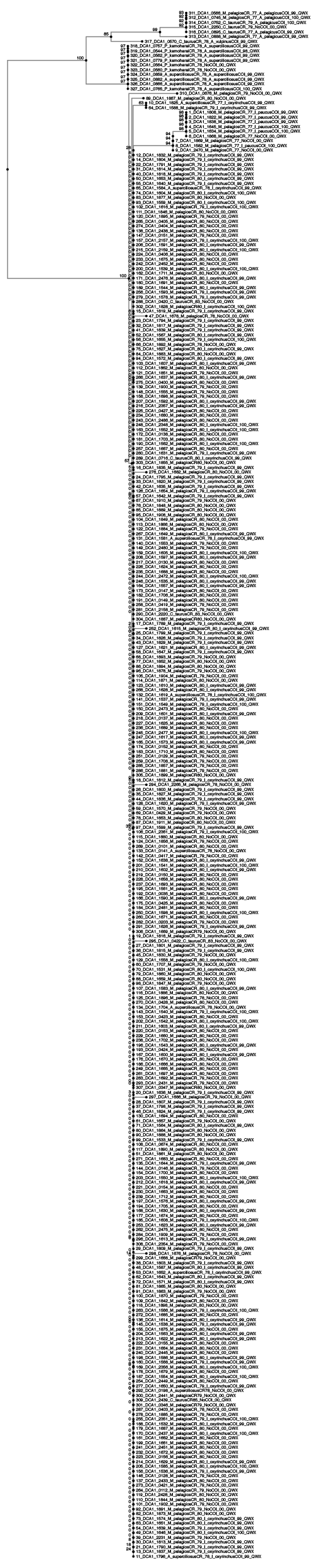
Numerous sequences amplified with CR primers from lamnid sharks that form a clade separate from the clade defined by vouchered whole mitochondrial genomes. These sequences have only a best match of 70-80% match to *M. pelagios* or *P. kamoharai* or *C. Taurus*, and group together with the putative pseudogene sequences in the upper clades of Figures 5 and 6. We use the species identification of a subset of these fins based on COI to define species clades, and use membership of a fin in one of these clades to assign the species name to that fin. For example, the first sequence at the top of this tree (DCA1_0566) came from a fin with a COI sequence with a 99% match to *A. pelagicus*. Another fin (DCA1_2250) is in this pseudo-CR sequence clade but has no COI sequence. We can use the presence of DCA1_2250 in this clade as good evidence that it is a fin from *A. pelagicus* as well.

#### Using pseudogenes in fin identifications

We can still use these pseudo-CR sequences in fin identification. First, we find fins that have been identified as *Isurus* or *Aolpias* species by their COI sequences. Then we use the CR sequences from these as references to identify species-specific clades of pseudo-CR sequences from other fins. Effectively, this uses the COI identifications as a reference set of fins from which we derive species-specific pseudo-CR data sets. This assignment identifies 305 sequences from *I. oxyrhincus*, 9 from *I. paucus*, 14 from *A. superciliosus*, and 8 from *A. pelagicus* (Fig. 7). These pseudogene numbers are in addition to the numbers of valid CR sequences: *I. oxyrhincus* =1, *I. paucus* =2, *A. superciliosus* =72, and *A. pelagicus* =0. However, an analysis of both data sets, (Fig. 8 see below), suggests caution in using CR sequences because species with these pseudogenes are seen in many fewer fins than suggested by the COI data set.

**Figure 8:**
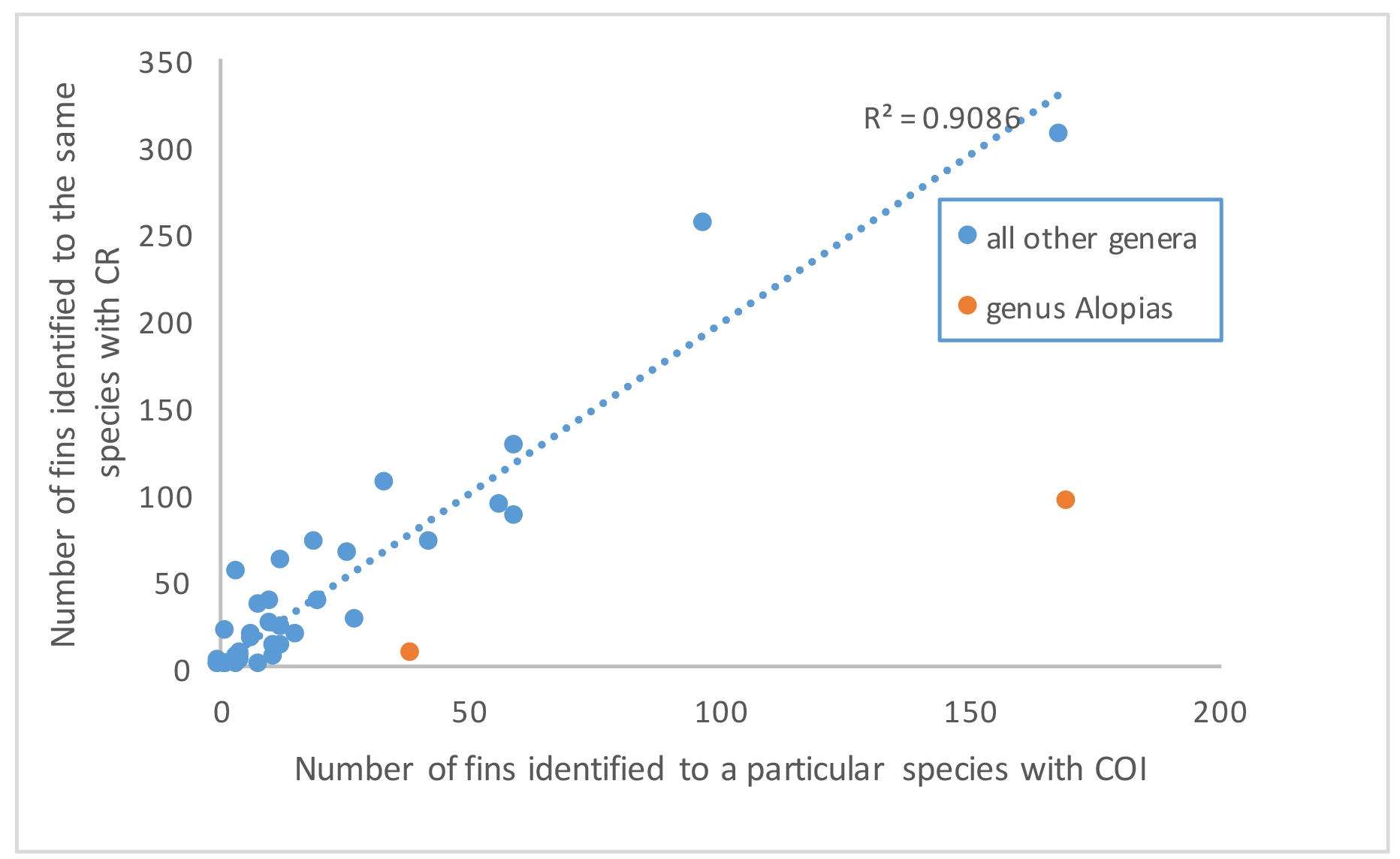
Good correlation between the number of fins seen for each species based on COI data (x-axis) compared to the control region (CR) data (y-axis). Two exceptions (red dots) are two species of Alopias, which appear to be underestimated with CR primers compared to COI primers.

#### Good CR amplifications for bacterial-laden samples

In our COI data set, 369 samples provided bacterial sequences. Of these, 284 amplified with our CR primers, and 236 provided good CR sequence, demonstrating that these samples retained shark mtDNA despite bacterial contamination. Most of these samples were from wet ceratotrichia collections that had been stored at room temperature for weeks to months before processing. This category of specimen is rare in shark fin samples. The species composition was similar to the composition from the rest of the CR data set. Six samples BLASTed to *Megachasma pelagios* at low similarity (80%), suggesting these were Lamnid sequences. Forty-eight sequences amplified the CR region but did not return good sequence.

#### Comparison of CR and COI identities

In our data set of 1528 CR sequences with strong matches to Genbank, we also have COI data for 802. Of these, 669 show a match between the species identities derived from the two different genes.

However, 133 show a high (>95%) match to a reference control region sequence but show a different species match (also at high % identity) for COI. This mismatch calls into question these sequence identifications. To resolve this, we re-sequenced a set of these mismatched samples for the mitochondrial 16S gene or cytochrome b and found that the 16S species identity matches the COI identity in about half the cases, and matches the CR identity in the other half. There are ten cases in which a third species is discovered this way. As a result of the uncertainty that this causes in species identifications, we have removed these sequences from our compilations.

#### Summary across all analyses

Table 2 compiles the species names derived from the CR data set, including cases where CR and COI labels match, when they do not match, and when we do not have COI data. We then show the species totals in Table 1, not including the cases where CR and COI sequences have high identity but do not match.

Overall, we can identify 969 fin products by COI sequence as belonging to 38-41 species, including 16-18 species of *Carcharinus*. We also identified 1,736 fins using control region sequences, showing 41-43 species, including the vast majority of the species from the COI data set. Table 1 shows that the most common species are thresher sharks, mako sharks, hammerhead and blue sharks, ocean species that have been seen widely in prior surveys of shark fin markets (e.g., Clarke et al., 2006). However, we also see a large number of samples and species of coastal requiem sharks in the genus *Carcharhinus*, as well as a diverse collection of hammerheads in seven of eight named species. The species targeted by the single biggest US shark fishery, spiny dogfish, does not appear among any of our fin products.

The species list in our shop data sets (summarized in Table 1) include 14-20 vulnerable species (IUCN Red list) and 3 endangered species. Six additional species the IUCN considers data deficient, and 6 species are specifically regulated under CITES. Altogether, the vulnerable, endangered and CITES species make up 36% of the species and 56% (CR) to 66% (COI) of the individual fin products. Of the CITES listed species, only the whale shark, basking shark and porbeagle (*Lamna nasus*) are not in our DNA sequence collection.

The Lamnid shark group and the hammerhead group have high fractions of vulnerable, endangered or CITEs regulated species, and both have high frequency in our sample. Likewise, seven of eight named hammerhead species are in our data set, including all five vulnerable, endangered or CITES regulated species. Only the newly described endemic *S. gilbert* is not in our data set.

Yet our data also show a wide variety of sharks that have not yet caught conservation attention, including a number of *Carcharhinus* species. We see 16-18 species of *Carcharhinus* sharks with an average number of individuals per species of 21 and 48 for COI and CR, respectively. These include widely ranging sharks such as the blacktip (*C. limbatus*), sandbar (*C. plumbeus*) and dusky sharks (*C. obscurus*). But we also see large numbers of more restricted species such as the shorttail shark (*C. porosus*) which lives in muddy estuaries in the western Atlantic, the blacknose (*C. acronotus*), inhabiting sandy habitats in the same regions, and the sharpnosed sharks (genus *Rhizopionodon*) found in Atlantic or Caribbean waters. Coastal sharks such as these are in contrast to the oceanic, pelagic sharks that have been the focus of a large shark fin fishery and large shark fin concern. Our data confirm that the diversity of coastal sharks is also a target of active shark finning.

## DISCUSSION

Our data show a large variety of sharks with great taxonomic diversity across the retail market sample. The COI data set provides identification of 36-39 species of sharks, including 17-19 *Carcharhinus* species, from 963 fin products. Likewise, the Control Region data set shows 38-41 species, including the vast majority of the species from the COI data set.

These species represent a large fraction of the known endangered, vulnerable or data deficient sharks in the world: more than 20 species in those categories occur in our data set. Moreover, 56-66% of the fins in our data set derive from these categories of sharks, largely because of the preponderance of thresher, mako and hammerhead sharks in the sample.

Our data bolster several recent retail market surveys that have also concluded that the shark fin market is highly diverse and highly endangered (Cardeñosa et al., 2018; Feitosa et al., 2018; Fields et al., 2018; Steinke et al., 2017). Steinke et al. (2017) found 20 species in 72 shark samples from Vancouver, Canada. Fields et al. (2018) found 59 species and 17 higher taxonomic groups when they monitored 3,952 samples from Hong Kong retail shops from 2014-2015. Similarly, Cardeñosa et al. (2018) found 82 species and species complexes in Hong Kong among small fin discards in 2016. Feitosa et al. (2018) found 17 predominantly nearshore species from 427 fin samples collected in northeastern Brazil. In these cases, and the samples reported here, some similarities emerge. All samples show a large number of endangered and threatened species. Thresher, hammerhead and mako sharks are abundant (especially *Alopias superciliosus, Sphyrna lewini*, and *Isurus oxyrhincus*). Abundant *Carcharhinus* samples include *C. falciformis, C. limbatus*, and *C. longimanus*, though there is a wide variety of other *Carcharhinus* species – generally at low and variable abundance - in the larger samples.

There are also some striking differences. The *Alopias/Isurus* group dominates the small Steinke sample (Table 1). The *Alopias/Isurus* group is also abundant in our sample, dominated by *A. superciliosus* and *I. oxyrhincus*. Yet in the Hong Kong shops sampled by Fields et al. (2018) and Cardeñosa et al. (2018), this group is a minor component. By contrast, the Fields et al. (2018) and Cardeñosa et al. (2018) samples are dominated by blue sharks, a single species in the Carcharhinidae that makes up nearly half their samples. Blue sharks occur in our sample, and in Steinke et al. (2017), at about 2-10%. Blue sharks also dominated the earlier sample of Clarke et al. (2006) from the Hong Kong wholesale market, prompting Fields et al. (2018) to ask how blue sharks could maintain their market presence at such a high level for so long. The relatively lower number of blue shark fins in retail markets on the US west coast may partially answer this question if there is a strong regional difference in market availability of this species. No blue sharks occur in the Brazil samples, which is dominated by regional sharpnose and *Carcharhinus* species (Feitosa et al., 2018).

### Comparison across genes and methods

There is strong evidence from these data that the CR and COI generally produce data equivalently well for most sharks, but that using CR primers specifically underestimates thresher shark abundance. The number of shark fins in our COI data set for each species is highly correlated to the number of fins we find with our CR primers (*r*^2^=0.91, Fig. 8). However there are two strong outliers. For both *Alopias superciliosus* and *A. pelagicus* the number of fins observed with CR primers, even taking in to account the fins identified with pseudo-gene CR sequences - falls far below the line relating COI and CR results (Fig. 8).

These outliers are because *Alopias* fins defined by COI sequences frequently fail to show *Alopias* CR sequences: 48 of the 208 *Alopias* fins with COI sequences have CR sequences with high identity to a different species, all *C. falciformis, C. obscurus*, and *C. longimanus* (see Table 3). These CR sequences are not *Alopias* pseudo-genes. Our hypothesis is that these CR identifications are incorrect, and derive from failure of the CR primers to easily amplify *Alopias* CR regions, amplifying common *Carcharhinus* species DNA as lab contaminants.

**Table 3:**
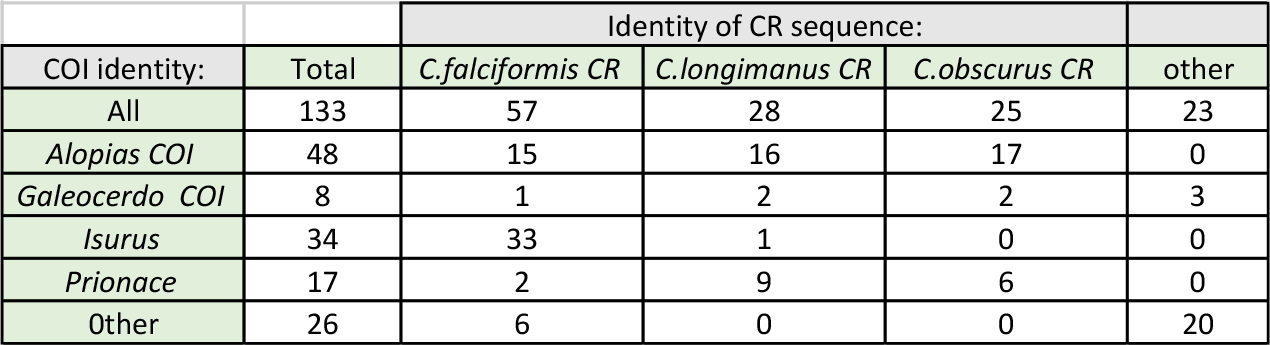
Fins with high fidelity COI sequence identifications (left column) that have different CR identifications, mostly to three species of common *Carcharhinus* sharks.

The lowest red point in Figure 8 represents very different identification rates for COI and CR for *A. pelagicus:* COI data show 38 fins for this species out of a total of 963 fins, but we find only 8 out of 1736 CR sequences. Lab records show that failure rate of CR amplifications of fins that have a *A. pelagicus* COI sequences is high. We searched for a possible explanation for this underrepresentation and found that the reverse primer from (Giles et al., 2016) does not match *A. pelagicus* at four of 22 positions, making it a poor amplification tool (Fig. 9). The match to *A. superciliosus* is better: 21 of 22 bases match. But the 3’ end base, the most crucial one for the polymerase chain reaction, does not match. This may make this primer less valuable for this species as well.

**Figure 9:**
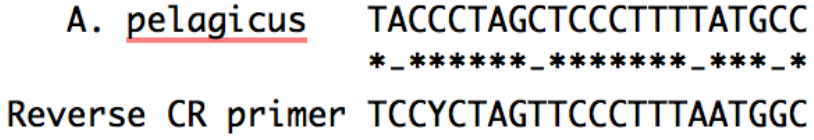
Mismatch between our reverse Control Region primer and the mitochondrial sequence of *Alopias pelagicus* may explain the strong underestimate of this species in our CR data set. The reverse CR primer is shown as the reverse complement of the synthesized reverse primer to correspond to the published sequence.

### Conservation status

Our data confirm the strong abundance of endangered, vulnerable and CITES listed shark species in retail fin markets. As in previous surveys, we find about half the fins come from species of conservation concern. This consistency is particularly important because it spans very different types of retail markets, in Hong Kong and the US west coast, imbedded in very different fisheries cultures and retail markets. In particular, the US has made strong strides in recent years in regulating fisheries, and returning overfished stocks to sustainability (Barner et al., 2015). Yet the shark markets in the US and Hong Kong remain quite similar in the presence of protected species. This suggests that fisheries policy improvements in the US have not penetrated the fin trade, and that highly regulated US fisheries do not contribute much to the retail fin market. This suggestion is supported by lack of fins from the biggest US shark fishery, spiny dogfish (*squalus acanthias*), both in the US west coast and Hong Kong retail fin market (Cardeñosa et al., 2018; Feitosa et al., 2018; Fields et al., 2018; Steinke et al., 2017).

There have been a series of other questions asked of the global shark fishery, notably about the persistence of some abundant fins on the global market from the same species for over a decade (Davidson, Krawchuk, & Dulvy, 2016; Fields et al., 2018). For example, blue sharks were a dominant feature of the Hong Kong market as shown in (Clarke et al., 2006), and again a decade later (Cardeñosa et al., 2018; Fields et al., 2018). Such dominance of a global market by a single species might suggest higher sustainability for this species than thought originally. However, our data show many fewer blue sharks, in contrast to nearly 50% of the samples being blue sharks in the Hong Kong retail market (Cardeñosa et al., 2018; Fields et al., 2018), we show no more that 4% in our survey. Likewise, bull sharks and sickle fin lemon sharks are at a much higher percentage in the Hong Kong market than in the US west coast. By contrast, our data show a higher abundance of bigeye thresher and the Atlantic smalltail sharks than seen in Hong Kong. These differences suggest that a single market can provide an incomplete picture of the global trade and call for more coordinated fin testing at high levels of sampling.

## ACKNOWLEDGMENTS

J. Giles provided assistance with data collection, technical issues, and laboratory supervision.

## AUTHOR CONTRIBUTIONS

SP oversaw data collection experiments, analyzed the data, contributed reagents/materials/analysis tools, and prepared figures and tables.

KR collected data and performed lab work.

SP, SJ, KV, KR conceived and designed the experiments, authored or reviewed drafts of the paper, and approved the final draft.

## DATA AVAILABILITY

All data and supplemental files are available open-access via the Open Science Framework, available at https://osf.io/9d65y/ and through the following DOI:10.17605/OSF.IO/9D65Y.

## SUPLLEMENTAL INFORMATION

Supplemental information for this article can be found online.

